# RNAtive to recognize native-like structure in a set of RNA 3D models

**DOI:** 10.1101/2025.07.01.662508

**Authors:** Jan Pielesiak, Maciej Antczak, Marta Szachniuk, Tomasz Zok

## Abstract

**Motivation:** Most widely used methods for evaluating RNA 3D structure models require experimental reference structures, which restricts their use for novel RNAs. They also often overlook recurrent structural features shared across multiple predictions of the same sequence. Although consensus approaches have proven effective in RNA sequence analysis and evolutionary studies, no existing tool applies these principles to evaluate ensembles of 3D models. This gap hampers the identification of native-like folds in computational predictions, particularly as AI-driven methods become increasingly prevalent.

**Results:** This paper presents RNAtive, the first computational tool to apply consensus-derived secondary structures for reference-free evaluation of RNA 3D models. RNAtive aggregates recurrent base-pairing and stacking interactions across ensembles of predicted 3D structures to construct a consensus secondary structure. It introduces a novel conditionally weighted consensus mode that treats interaction networks as fuzzy sets and uniquely allows integration of user-defined 2D structural constraints, enabling evaluation guided by experimental data. Input RNA models are ranked using two adapted binary-classification-based scores. Benchmarking against CASP15 competition data shows that models consistent with the consensus exhibit native-like structural features. The RNAtive web server offers an intuitive platform for comparing and prioritizing RNA 3D predictions, providing a scalable solution to address the variability inherent in deep learning and fragment-assembly methods. By bridging consensus principles with 3D structural analysis, RNAtive advances the exploration of RNA conformational landscapes and has potential applications in fields like therapeutic RNA design.

**Availability:** RNAtive is a freely accessible web server with a modern, user-friendly interface, available for scientific, educational, and commercial use at https://rnative.cs.put.poznan.pl/.

## 1 Introduction

RNA molecules exhibit remarkable structural versatility, adopting diverse three-dimensional (3D) conformations influenced by environmental factors and molecular interactions [Haque et al., 2017, Schroeder et al., 2004, Vicens and Kieft, 2022, Leamy et al., 2016, Pai and Luca, 2018]. While a single sequence can yield multiple stable states, as in riboswitches and aptamers, evolutionary pressure preserves a functional structural core [Bevilacqua and Tolbert, 2022]. These conserved core motifs – such as catalytic sites or ligand-binding pockets – remain invariant across diverse conformations, while peripheral regions may vary. Identifying these recurrent motifs is central to the concept of consensus, which enables the extraction of biologically meaningful patterns from structural diversity [Haque et al., 2017, Leamy et al., 2016].

In bioinformatics, consensus approaches are robust strategies for extracting signals from heterogeneous data. They are used to identify functional motifs in sequence alignments, infer conserved base-pairing patterns from multiple sequence alignments to improve RNA secondary structure prediction, and define invariant cores from ensembles of experimental 3D models [Hofacker, 2007, Bernhart and Hofacker, 2009, Hamada, 2014, Fukunaga and Hamada, 2022, Pietrosanto et al., 2016, Zhao and Wang, 2008].

Consensus principles, however, remain underutilized in the assessment of computational RNA 3D structure predictions. Current evaluation strategies predominantly rely on reference-dependent metrics, such as Root Mean Square Deviation (RMSD) or Interaction Network Fidelity (INF), that require an experimental structure, limiting their applicability to novel RNAs [Parisien et al., 2009, Zok et al., 2013, Wiedemann et al., 2017, Magnus et al., 2019, Mackowiak et al., 2024]. Moreover, they fail to capture recurrent structural features across predicted conformations – a critical shortcoming given the high variability of models from emerging deep-learning methods. While other approaches can flag outliers or predict model quality, they do not systematically extract consensus signals from ensembles of predicted structures [Davis et al., 2007, Luwanski et al., 2022, Townshend et al., 2021].

To address this gap, we introduce RNAtive, a web-based tool that establishes a new paradigm for evaluating ensembles of RNA 3D models. Unlike traditional methods that score models in isolation, RNAtive exploits the synergistic information within the entire set of models to identify the most plausible structures when no reference is available. Beyond this conceptual shift, RNAtive incorporates several algorithmic innovations. The conditionally weighted consensus mode introduces a novel, threshold-free approach that treats interaction networks as fuzzy sets. Accordingly, both INF and the F1-score were adapted to operate within this fuzzy-set framework. Furthermore, RNAtive is the first tool to support the integration of 2D structural constraints into the consensus calculation, a vital capability for guiding model evaluation with prior experimental data. Together, this combination of a consensus-driven paradigm, a fuzzy-set algorithmic framework, and constraint integration constitutes a significant contribution that enables analyses previously impossible to perform in an automated way.

## 2 System and methods

### 2.1 Input and output

RNAtive accepts tertiary structures in both PDB and PDBx/mmCIF formats. Users can upload multi-model files or ZIP/TAR.GZ archives, which are automatically unpacked and divided into individual models. Because RNAtive relies on nucleotide interactions, all input models must be consistent regarding nucleotide composition. If needed, RNAtive remaps chain names and renumbers residues internally to achieve the expected consistency. However, all other differences in nucleotide composition cannot be automatically resolved. Thus, they are detected and prevented from submitting the inconsistent model.

As an optional pre-processing step for practical data sanitation, RNAtive incorporates an embedded MolProbity filter [Davis et al., 2007]. This feature is conceptually distinct from the core ranking algorithm: whereas the primary methodology is fundamentally consensus-driven, deriving its signal from the internal agreement within an ensemble, the filter serves as a knowledge-based heuristic to identify and exclude models with severe stereochemical flaws that could otherwise distort the consensus calculation. Its application is entirely at the user’s discretion.

The filter utilizes MolProbity’s classification of structural features, which assigns models a label of *good, caution*, or *warning* based on criteria such as clashscore (number of atomic clashes per 1,000 atoms) as well as occurrences of bad bonds or bad angles. Users may then exclude models with high clashscores or other stereochemical warnings prior to consensus ranking.

Optionally, the entire analysis can be subjected to 2D structure constraints. Users can provide a canonical 2D structure in dot-bracket format, using brackets (), [], etc. for expected base pairs, dots. for unconstrained nucleotides, and x to indicate that a nucleotide should be unpaired (i.e., not forming either canonical or non-canonical base pairs). The inclusion of x allows dots to be interpreted as nucleotides that may form non-canonical interactions, consistent with how canonical 2D structures are typically encoded in dot-bracket format. This functionality, unique to RNAtive, allows users to guide the evaluation with prior experimental knowledge – a critical feature for practical applications that no other method provides.

Users can also configure the analysis by selecting one of six available tools for RNA interaction annotation: RNApolis Annotator [Szachniuk, 2019], BPNet [Roy and Bhattacharyya, 2022], FR3D [Sarver et al., 2007], MC-Annotate [Gendron et al., 2001], RNAView [Yang, 2003], or Barnaba [Bottaro et al., 2018]. Additionally, RNAtive supports two modes of operation. If conditionally weighted consensus is enabled, RNAtive treats annotated interactions as a fuzzy set, where an interaction’s membership is determined by its frequency of occurrence across all analyzed models. Alternatively, RNAtive can be parameterized with a confidence level, which defines a minimum frequency threshold for an interaction to be considered.

The results page provides an overview of the input parameter values and constraints, the derived consensus (including visualization and interaction tables), final model rankings, and detailed information for each model, including its MolProbity quality scores, if computed.

### 2.2 RNAtive workflow

The RNAtive workflow (Figure 1) is organized into three main stages: data preparation, consensus construction, and scoring with ranking, which are described in the following sections.

**Figure 1:**
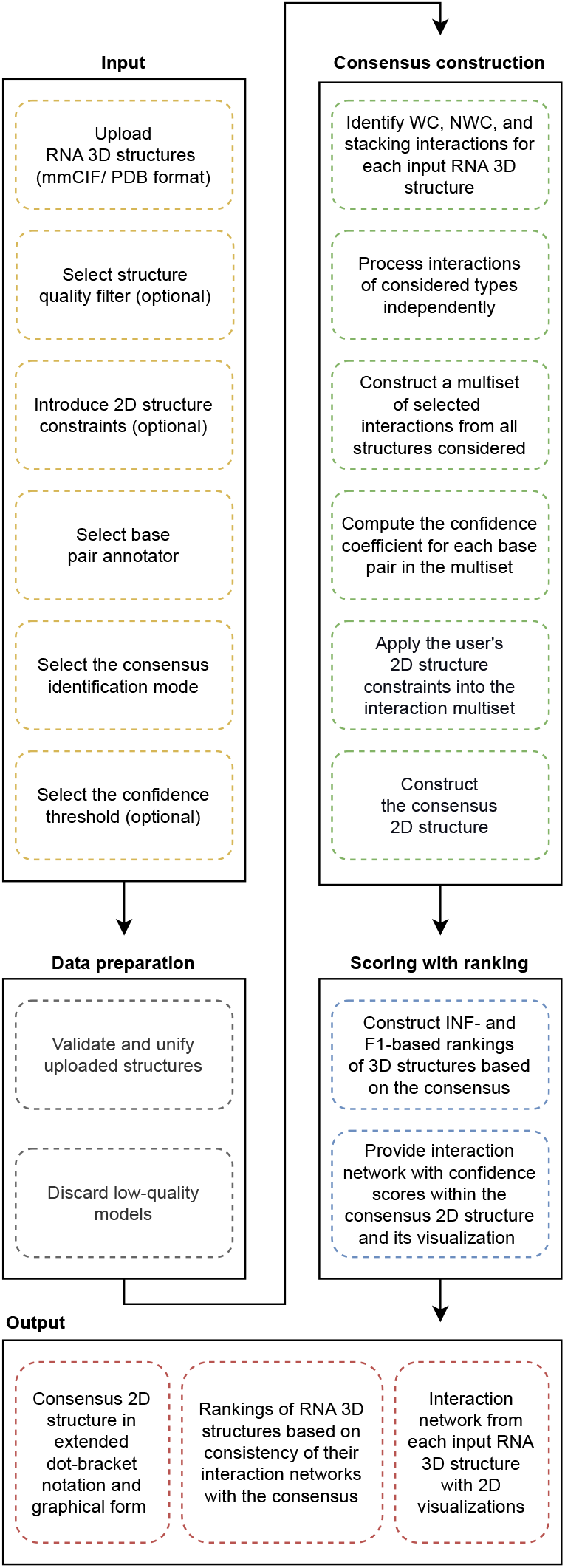
RNAtive workflow.

#### Data preparation

RNAtive starts with a unification protocol to ensure consistent chain names and residue numbering in submitted RNA 3D structures (Figure 1). All inputs are converted to PDB format since some external software, including MolProbity, only supports this format. It is essential to note that the conversion from PDBx/mmCIF to PDB can fail under certain conditions: 1) if the atom count exceeds 99,999; 2) if the residue count exceeds 9,999; or 3) if the chain count exceeds 62, which includes a combination of uppercase letters, lowercase letters, and digits. We recognize these limitations of RNAtive. However, for its primary purpose – analyzing computationally derived RNA 3D models – we believe the system can effectively handle any realistic dataset.

Once the model consistency is ensured, MolProbity (if selected by the user) filters out models that do not meet the user’s quality expectations. The processing continues only if at least two models pass this stage.

#### Consensus construction

All valid models proceed to interaction analysis using the chosen annotation tool. This analysis produces a set of unique interactions – base pairs and stacking – each accompanied by metadata describing its frequency among the models and flags indicating compliance with or violations of the provided 2D constraints. We treat this multiset of interactions and metadata as the virtual consensus structure representing the ensemble of input RNA 3D models (Figure 2). A similar interaction set with metadata is also computed for each model individually for scoring.

**Figure 2:**
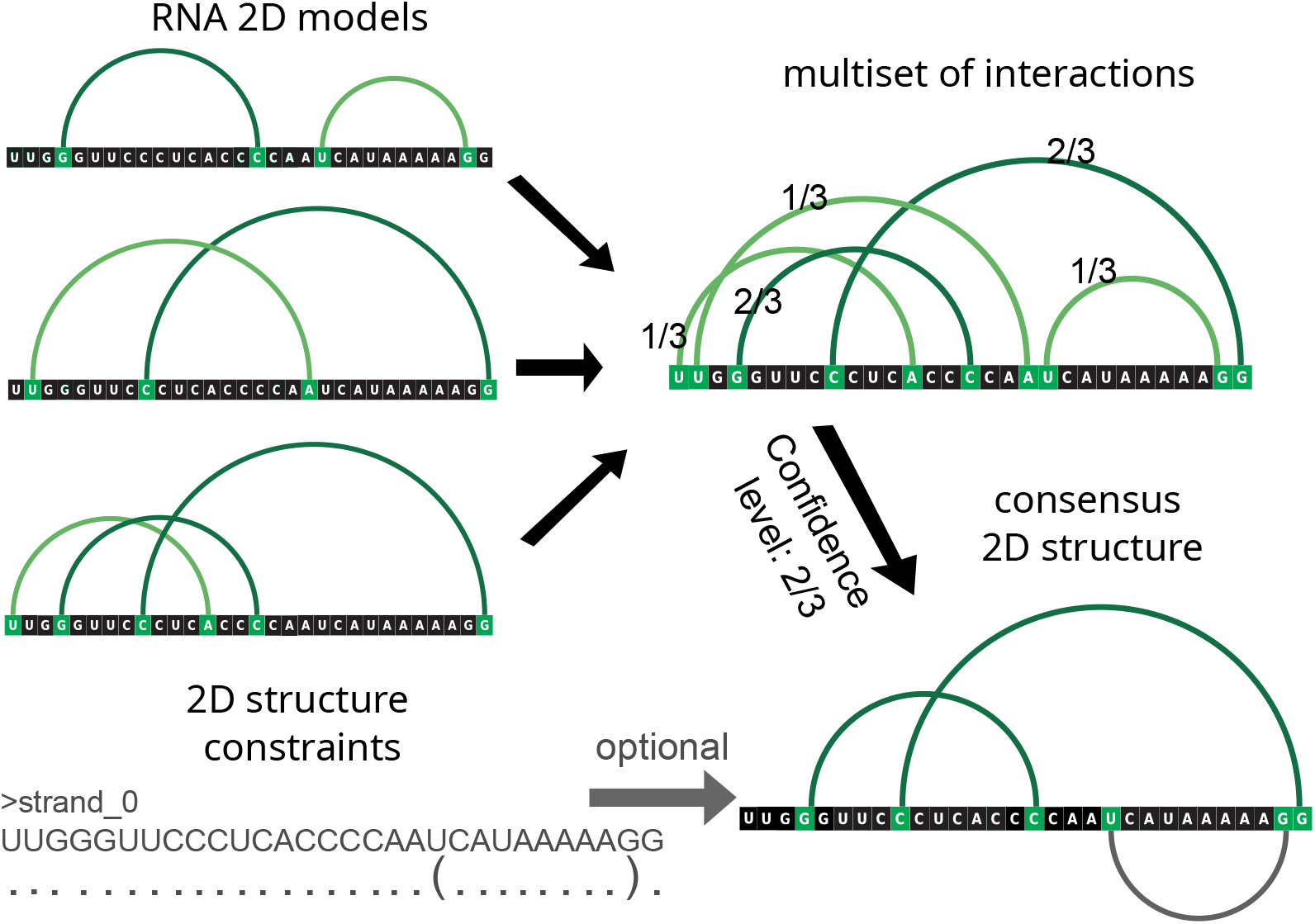
Example secondary structure models derived from input 3D structures, the resulting multiset with base pair frequencies, and the consensus secondary structure. The additional constraint (bottom left) favors the first model and penalizes the other two, even though the constrained base pair (grey) is excluded from the consensus at the selected confidence level of 2/3.

The consensus structure depends on user input and can be constructed in two modes: conditionally weighted or threshold-based. In the conditionally weighted mode, interaction sets are treated as fuzzy sets, where frequencies are directly incorporated. This approach weights common interactions more heavily, rewarding models that include them and penalizing those that omit them. In the threshold-based mode, a consensus is defined by selecting a confidence level *n* (ranging from 1 to the number of input models). Only interactions observed in at least *n* models are retained. Each model is then compared against the resulting consensus structure. Thus, users can choose between a continuous, frequency-weighted consensus and a discrete, threshold-based consensus, depending on their specific needs.

#### Scoring and ranking

The interaction sets are used to compute RNAtive outputs. Two primary scores are reported for each RNA 3D model: INF and the F1-score. In the threshold-based mode (when the confidence level parameter is set), the following definitions apply: True Positives (*TP*) are interactions found in both the model and the consensus subset; False Positives (*FP*) are interactions present only in the model; and False Negatives (*FN*) are those found only in the consensus. INF and F1-score are then computed according to Eq. 1 and 2, respectively.

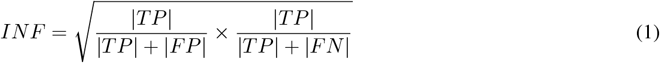

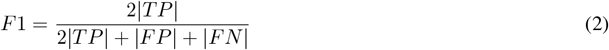

When the conditionally weighted consensus mode is active, we adapt the standard definitions of INF and F1-score. Specifically, if an interaction with consensus membership *p* is present in both the consensus and a given model’s set, *p* is added to *TP*_*sum*_, and 1 − *p* is added to *FP*_*sum*_. If the interaction is in the consensus but absent from the model, *p* is added to *FN*_*sum*_. With this scheme, uncommon consensus interactions (*p <* 0.5) contribute significantly to *FP*_*sum*_ when included in the model, but only weakly to *FN*_*sum*_ when omitted. Conversely, common interactions (*p >* 0.5) increase the similarity score for models that include them and decrease it for models that do not. This approach provides a nuanced use of all interaction data from RNA 3D models without requiring a predefined confidence level, making it particularly relevant for consensus-based analyses. The formulas for INF and F1-score remain unchanged (Eq. 1 and 2), except that the absolute counts |*TP*|, |*FP*|, and |*FN*| are replaced by the corresponding sums *TP*_*sum*_, *FP*_*sum*_, and *FN* _*sum*_

### 2.3 Implementation

The RNAtive application consists of multiple components orchestrated using Docker Compose. The frontend is built with the React framework and the Ant Design UI library, while the backend is implemented in Java 17 using the Spring framework. The system also integrates a PostgreSQL database engine and Nginx, the gateway and ingress controller. RNAtive incorporates the RNApdbee 3 adapters container [Zok et al., 2018], which provides base pair annotation and visualization tools. It also uses the cli2rest-rnapolis, cli2rest-varna-tz, and cli2rest-rchie Docker containers for structural unification and customized visualization via VARNA and R-Chie [Darty et al., 2009, Tsybulskyi et al., 2020]. The web application is served over HTTPS with HAProxy – a high-performance load-balancer – and secured using Let’s Encrypt certificates managed by Certbot automation [Aas et al., 2019].

### 2.4 Performance evaluation methodology

We evaluated RNAtive by comparing it with other state-of-the-art approaches, which can be broadly categorized into knowledge-based and machine learning (ML)-based methods. The knowledge-based techniques, which are more numerous, rely on calculating statistics-driven scores for atomic systems and assessing models accordingly. They differ mainly in how the scoring is performed and which factors are considered. In contrast, ML-based methods employ diverse architectures to rank RNA 3D models. Further details about these methods are provided in Table 1.

**Table 1:** State-of-the-art approaches for RNA 3D model quality assessment.

| Method | Reference | Description |
| --- | --- | --- |
| **Knowledge-based approaches** |  |  |
| RASP | Capriotti et al. [2011] | An all-atom statistical potential to rank models. |
| 3dRNAscore | Wang et al. [2015] | Distance- and dihedral-dependent energies to evaluate models. |
| DFIRE | Zhang et al. [2020] | All-atom distance-dependent energy function based on a reference state (distance-scaled finite ideal-gas reference state). |
| BRiQ | Xiong et al. [2021] | Statistical energy function with a nucleobase-centric sampling algorithm, capturing full-orientation dependence of base-base, oxygen-base, and oxygen-oxygen interactions with the RNA backbone. Quantum-mechanical calculations ensure proper weights to minimize the possible effects of indirect interaction. |
| rsRNASP | Tan et al. [2022] | All-atom distance-dependent statistical potential based on residue separation. |
| cgRNASP, cgRNASP-C, cgRNASP-CN, cgRNASP-PC | Song et al. [2022],<br>Tan et al. [2023] | Direct successor of rsRNASP, uses both long- and short-range interactions by residue separation. |
| **Machine learning-based approaches** |  |  |
| RNA3DCNN_MD, RNA3DCNN_MDMC | Li et al. [2018] | Uses a Convolutional Neural Network to analyze every nucleotide individually with a 3D grid representation without manual structure extraction. |
| ARES | Townshend et al. [2021] | A deep neural network with an architecture formulated for direct 3D structure input that predicts the model’s RMSD using the spatial position of atoms and their chemical element types. |
| PAMNet | Zhang et al. [2023] | The Physics-Aware Multiplex Graph Neural Network that induces a physics-informed bias that allows for modeling of local and non-local interactions of molecules based on different geometric information. |
| lociPARSE | Tarafder and Bhattacharya [2024] | A Locality-aware invariant point attention architecture that estimates the IDDT scores to assess the accuracy of each nucleotide and the atomic environment that surrounds it in a superposition-free manner. |

In the benchmark, we used two RNA 3D structure datasets: CASP15 [Kryshtafovych et al., 2023, Das et al., 2023] and the randstr decoy set [Capriotti et al., 2011]. Both contain experimentally determined target structures, each associated with an ensemble of 3D models, but their origins differ. The CASP15 set comprises diverse predictions submitted by independent groups modeling structures from a given sequence, whereas the randstr set contains more uniform decoys generated computationally by perturbing native structures. The CASP15 dataset includes 12 targets (1,672 models), while the randstr dataset comprises 85 targets (42,585 models).

To establish a non-redundant set of structural quality metrics for evaluating 3D models, we systematically analyzed correlations among them. Each model in the CASP15 and randstr datasets was assigned multiple scores used for global ranking.

The CASP15 dataset provides pre-computed scores for RMSD, local Distance Difference Test (lDDT; Mariani et al. [2013]), Template Modeling score (TM-score; Gong et al. [2019]), and Global Distance Test–Total Score (GDT-TS; Zemla [2003]). For the randstr dataset, the available scores include RMSD-C3’, GDT-C3’, RMSD-ALL, and GDT-ALL. To supplement these, we calculated three INF-based scores for every model: INF_WC_ (for canonical base pairs), INF_nWC_ (for non-canonical base pairs), and INF_all_ (for all base pairs) [Parisien et al., 2009]. These scores were computed from nucleotide interactions annotated using FR3D [Sarver et al., 2007].

Our objective was to remove structural quality metrics whose rankings were highly correlated. To identify such redundancy, we computed Spearman’s *ρ* correlation matrix across all scores within each dataset. We then used Fisher’s z-transformation to average the correlation coefficients across the 12 CASP15 targets and the 85 randstr targets. For every pair of metrics with opposite directions (e.g., RMSD as a dissimilarity measure versus lDDT as a similarity measure), we inverted the sign of the correlation coefficient. Finally, the correlation matrix was transformed into a dissimilarity matrix and subjected to hierarchical clustering using the Unweighted Pair Group Method with Arithmetic mean (UPGMA).

Cutting the resulting dendrograms (see Figures S1–S2 in the Supplementary data) at a dissimilarity threshold of 0.25 revealed distinct, densely connected clusters. For the CASP15 dataset, this analysis identified four groups: one cluster containing INF_WC_, INF_all_, and lDDT; a second cluster with TM-score and GDT-TS; and two single-element clusters for RMSD and INF_nWC_. Based on these results, we selected four representative structural quality metrics for further analysis: INF_nWC_, RMSD, TM-score, and lDDT. TM-score was chosen as a modern alternative to the highly correlated GDT-TS, and lDDT as a widely used measure of local similarity.

For the randstr decoys, clustering revealed that RMSD-ALL, RMSD-C3’, GDT-ALL, and GDT-C3’ are highly redundant and form a single large cluster. A second cluster contained INF_WC_ and INF_all_, while INF_nWC_ again formed a single-element cluster. Consequently, we proceeded with three metrics: INF_nWC_, RMSD-ALL, and INF_WC_. RMSD-ALL was selected as a widely recognized global metric encompassing all atoms, and INF_WC_ was preferred over INF_all_ to avoid data dependency, since INF_all_ overlaps with the already selected INF_nWC_.

Next, we applied a suite of state-of-the-art evaluation tools (Table 1) to score each model’s PDB file. This process generated a quality ranking for the models of each target according to each tool. We then compared these tool-generated rankings against reference rankings derived from the selected structural metrics (four for CASP15 and three for randstr, as described above).

To quantify the concordance between tool- and reference-based rankings, we selected four distinct ranking similarity measures:

- Spearman’s *ρ*, a standard measure of rank correlation.
- Kendall’s *τ*, which counts the number of concordant and discordant pairs between two rankings.
- Enrichment Score (ES) [Tsai et al., 2003], a top-k overlap metric commonly used in structural bioinformatics.
- Rank-Biased Overlap (RBO) [Webber et al., 2010], which compares entire rankings while giving greater weight to agreement on top-ranked items.

A critical step in this comparison was to harmonize the scoring directions. Some tools output scores where lower values indicate better performance (e.g., energy), whereas reference metrics sometimes score higher as better (e.g., lDDT). We accounted for this by inverting the signs of the correlation coefficients or sorting one of the rankings in reverse order as needed.

Although the four selected measures are distinct, they are inherently correlated, as they all assess ranking similarity. To synthesize them into a single robust score, we applied Principal Component Analysis (PCA) and extracted the first principal component (PC1). Before PCA, we transformed the data to stabilize variance and meet model assumptions: values in the [−1, 1] range (Spearman’s *ρ* and Kendall’s *τ*) were transformed using the inverse hyperbolic tangent (arctanh), and values in the [0, 1] range (ES and RBO) were transformed using the logit function. The resulting values were then standardized.

Our analysis confirmed the validity of this approach. All PC1 loadings were positive, indicating that the scoring directions were correctly aligned across tools and metrics. Moreover, PC1 alone explained 76% and 80% of the variance in the CASP15 and randstr datasets, respectively. This high explanatory power demonstrates that the four metrics capture a common underlying concept of ranking quality and that PC1 serves as an effective composite score for how well a tool’s ranking aligns with a reference standard for a given target.

Using this unified PC1 score, we carried out two analyses. First, we generated detailed per-metric ranking plots to visualize the performance of each tool individually. Second, we conducted a global statistical comparison. We began with Friedman’s test, which confirmed significant performance differences among the tools (p-values were near zero for both datasets). This allowed us to proceed with a Nemenyi post-hoc test to perform all pairwise comparisons and identify which tools differed significantly. Finally, we summarized the results using Critical Difference (CD) diagrams, which provide a global performance ranking and visually cluster tools whose performances are not statistically distinguishable.

### 2.5 Robustness analysis methodology

To rigorously evaluate the robustness of RNAtive, we performed statistical tests using a noise-injection simulation protocol. The analysis employed a dataset derived from 13 structural models of the thymidylate synthase mRNA regulatory motif, the first challenge of RNA-Puzzles. RNAtive was run with default settings: no quality filtering, no 2D structure constraints, RNApolis Annotator [Szachniuk, 2019] for base pair analysis, and a conditionally weighted consensus. We exported all 840 canonical and non-canonical base pairs and stacking interactions, comprising 116 unique interaction types. This dataset served as the input for a custom simulator designed for the robustness analysis.

The simulator introduced systematic, random noise into the dataset iteratively. In each iteration, it randomly selected an RNA 3D model to alter, an interaction type (canonical, non-canonical, or stacking), and an action (“add” or “remove”). For an “add” operation, the simulator randomly generated a new interaction – a pair of nucleotide identifiers for canonical or stacking interactions, or a triplet including Leontis-Westhof classification for non-canonical interactions – continuing until a previously absent entry was found. For a “remove” operation, an existing interaction was randomly selected and removed.

First, a single reference ranking was generated by running RNAtive on the original, unaltered dataset. Subsequently, after each iterative alteration, RNAtive was re-executed to generate a new noisy ranking. We ran 100 independent simulations, each with 1,000 iterations. This process yielded, for each noise level *n* (where 1 ≤ *n* ≤ 1000), a sample distribution of 100 rankings derived from *n* random alterations, which were then compared to the baseline.

Performance was evaluated by comparing each noisy ranking to the reference ranking using four similarity metrics: Spearman’s *ρ*, Kendall’s *τ*, ES, and RBO. The standard ES definition [Wang et al., 2015] considers the top 10% of results, which in this case (13 models) corresponds to the single top-ranked position. To provide a more informative assessment, we adapted the metric to focus on the top 25% (i.e., the first three positions), denoted ES_25%_ to distinguish it from the standard ES used elsewhere in the manuscript.

To establish a performance baseline, we constructed a null model representing a random ranker. We generated 10,000 batches, each containing 100 random pairs of 13-item rankings. We then calculated the mean for each of the four similarity metrics separately across these 100 pairs. Repeating this for all 10,000 batches produced four distinct sampling distributions of mean values (one for each metric), from which the 95% confidence intervals (CIs) were derived. The null-model analysis revealed 95% CIs of [− 0.057, 0.057] for Spearman’s *ρ* and [− 0.041, 0.041] for Kendall’s *τ*. The 95% CI for ES_25%_ was [1.87, 2.73], and the RBO 95% CI was [0.522, 0.555].

## 3 Results and Discussion

### 3.1 Performance on the CASP15 dataset

The performance of RNAtive was first evaluated with INF_nWC_ (Figure 3A), chosen as a reference metric due to its low correlation with other structural quality measures. On the CASP15 models, RNAtive variants showed clear dominance, occupying the top nine positions as well as the 11th and 12th. This indicates that RNAtive’s core algorithms are highly effective at addressing the structural challenges presented in CASP15. However, no consistent advantage was observed between the “all” and “canonical” interaction models. The top two methods – *RNAtive RNAView all* and *RNAtive FR3D all* – are closely followed by *RNAtive BPNet canonical, RNAtive RNApolis canonical*, and *RNAtive FR3D canonical*. Interestingly, these canonical-only variants outperformed their “all” counterparts (e.g., *RNAtive BPNet all* ranked 7th), suggesting that, for CASP15 targets, the inclusion of non-canonical interactions provides inconsistent benefits. This may reflect either a limited role of such interactions in model ranking or reduced accuracy of annotation tools in detecting non-canonical pairs, with the latter potentially introducing noise that degrades overall ranking.

**Figure 3:**
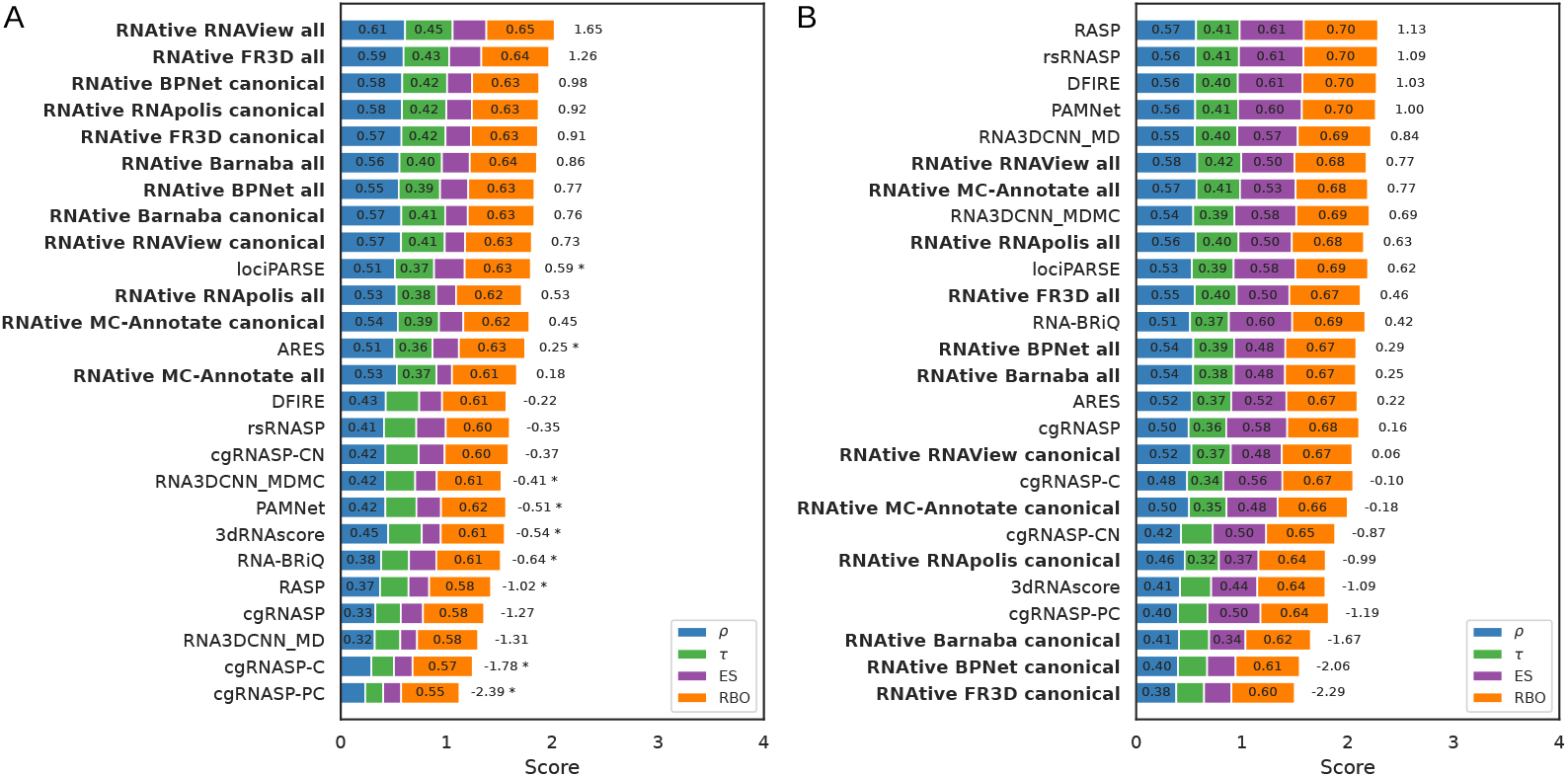
Ranking performance of scoring methods for the (A) CASP15 and (B) randstr decoy datasets. Each plot evaluates how the scoring methods rank RNA 3D models relative to the INF_nWC_ structural quality metric. Each bar reports four ranking similarity measures, averaged across all targets: Spearman’s *ρ*, Kendall’s *τ*, the normalized ES, and RBO. Methods are ordered by their score on the PC1 from the PCA of these four measures, providing a unified performance assessment. The PC1 score is labeled on each bar; its magnitude determines the ranking, though the absolute value is not directly interpretable. An asterisk (*) indicates that for a given method, ≥ 20% of the underlying correlation coefficients (*ρ* or *τ*) were not statistically significant. Results for RNAtive, tested under different parameter settings, are shown in bold.

The overall performance across all reference metrics for the CASP15 dataset is summarized in a Critical Difference (CD) diagram (Figure 4A). The diagram reveals a clear hierarchy led by the RNAtive family of tools. Within this dominant group, an elite tier emerges: the top two performers, *RNAtive FR3D all* and *RNAtive RNAView all*, show statistical superiority over a broader set of competitors than any other method. This highlights the exceptional effectiveness of RNAtive’s core algorithms, particularly these two variants, in addressing CASP15-like challenges.

**Figure 4:**
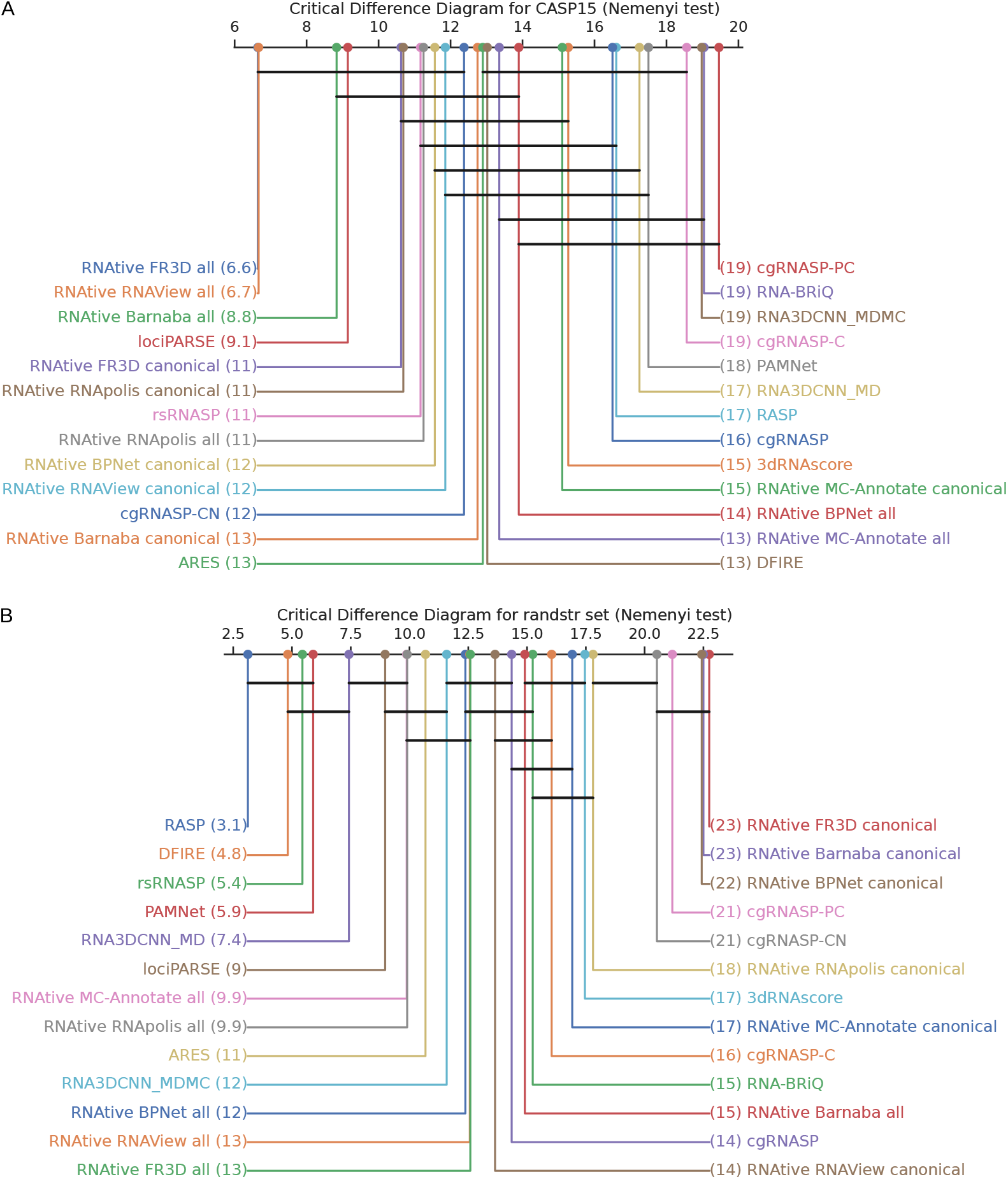
CD diagram for tool performance on the (A) CASP15 and (B) randstr decoys. The diagram shows the average rank of each tool across all evaluation tasks. Tools are ranked from best (left) to worst (right). Horizontal bars connect groups of tools whose performance is not statistically different according to a post-hoc Nemenyi test (*p <* 0.05).

### 3.2 Performance on the randstr dataset

In contrast to the CASP15 results, the performance of the RNAtive family declines markedly on the randstr dataset (Figure 3B). The best-performing variant, *RNAtive RNAView all*, ranks only 6th overall, with other “all” variants clustered in mid-range positions. Even more striking is the weak performance of the “canonical” variants. A pronounced split emerges: all RNAtive “all” variants fall within the top half of the ranking, whereas every “canonical” variant is confined to the bottom half.

The CD diagram for the randstr dataset (Figure 4B) shows a complete reversal of the performance hierarchy. A new top tier – comprising RASP, DFIRE, rsRNASP, and PAMNet – now significantly outperforms the entire RNAtive family. The RNAtive “all” variants are relegated to a crowded middle tier, alongside methods such as lociPARSE and ARES. Notably, several of the new leaders (e.g., RASP, DFIRE, and PAMNet) had previously ranked in the lowest tiers on the CASP15, underscoring their specialization for the randstr task.

### 3.3 Dataset-specific strengths revealed in benchmarking

Taken together, our analysis demonstrates that scoring tool performance is highly dataset-dependent and reveals a crucial trade-off between peak performance on a specialized task and robust generalization. The behavior of RASP exemplifies this trade-off: its Spearman’s *ρ* (against INF_nWC_) falls dramatically from a strong 0.57 on the randstr to a modest 0.37 on CASP15. In stark contrast, leading RNAtive variants like *RNAtive RNAView all* demonstrate remarkable consistency, maintaining high Spearman’s *ρ* scores of 0.61 on CASP15 and 0.58 on the randstr set. Meanwhile, other tools, such as lociPARSE, prove to be consistent mid-tier performers across both datasets. This suggests that energy-based scoring methods may be highly tuned for decoy sets, where models are often subtle perturbations of one another, while RNAtive’s geometry-based consensus approach proves more stable when faced with the heterogeneous ensembles typical of a CASP experiment.

This stability highlights RNAtive’s main advantages: its robust, interpretable, and highly flexible performance. While many state-of-the-art methods operate as rigid, pre-trained models that struggle to generalize, RNAtive maintains stable performance across diverse datasets. This flexibility is twofold. First, RNAtive is intentionally designed to be agnostic to the source of the input models, empowering users to apply our tool to any ensemble, whether generated by a single method, a combination of established tools, or novel prediction algorithms. Second, its unique configurability enhances this robustness, allowing users to guide the analysis by applying quality filters, introducing 2D structure constraints, and fine-tuning the consensus definition with confidence thresholds or conditional weighting – features other methods lack. Crucially, RNAtive is not a black box as it presents the consensus structure it derives, providing direct insight into the recurrent interaction patterns that underpin its high-quality results.

### 3.4 Robustness to noise in input data

The robustness of RNAtive was evaluated using the noise-injection protocol described in Section 2.5. Despite increasing levels of random perturbation, RNAtive consistently maintained strong performance. As shown in Figure 5, Spearman’s *ρ* declines with added noise, but the mean remains around 0.6 even at the highest noise level – well above the null-model’s 95% CI, which peaks at 0.057. Comparable stability is observed for Kendall’s *τ*, ES_25%_, and RBO (see Figures S8–S10 in the Supplementary data).

**Figure 5:**
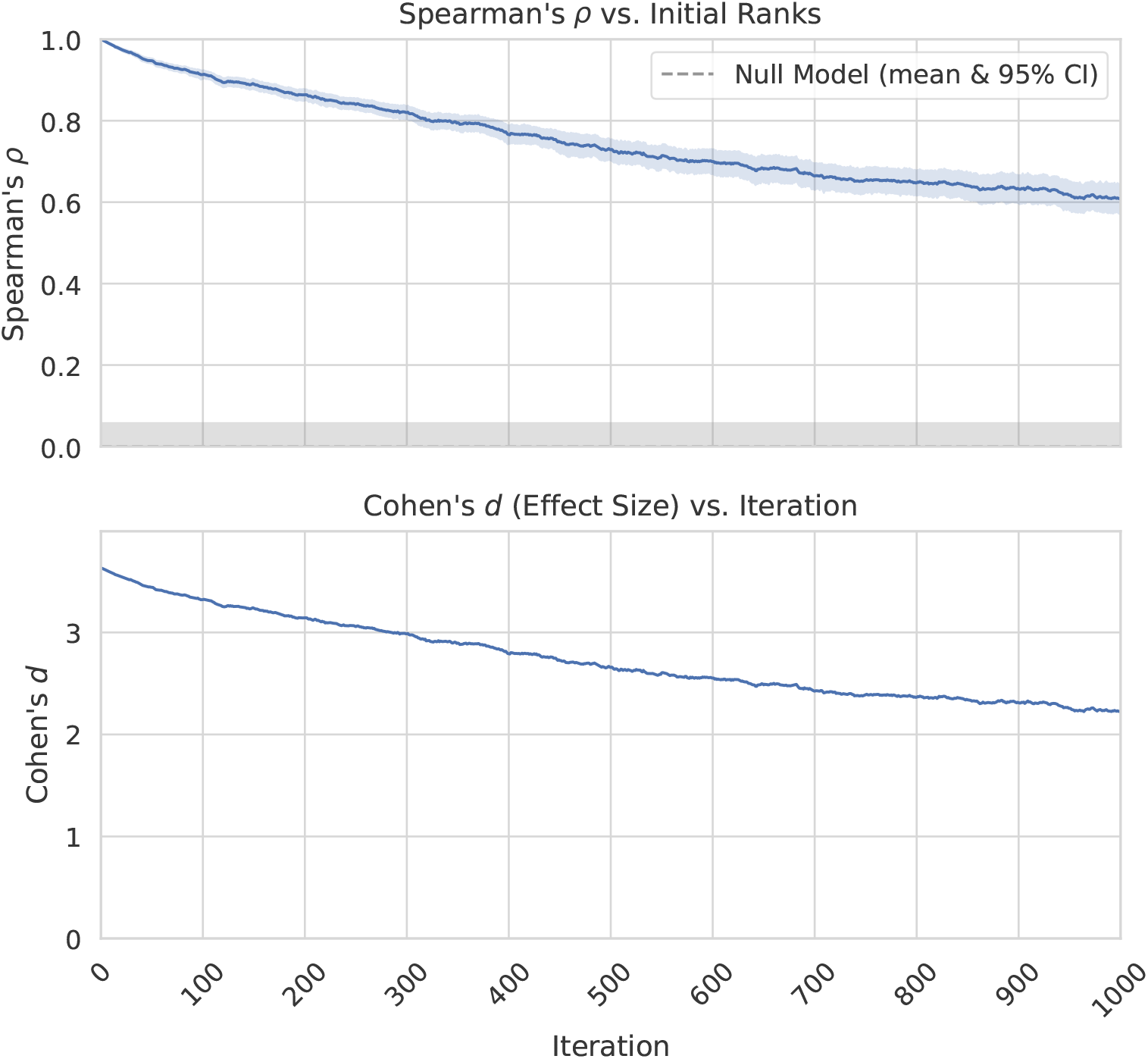
The effect of increasing levels of random noise on RNAtive performance. At the top, Spearman’s correlation coefficient (*ρ*), shown with a 95% confidence interval (CI), remains clearly distinct from the null-model’s mean and 95% CI, even at the maximum applied noise level. At the bottom, the effect size, quantified by Cohen’s *d*, remains large across the entire simulation, confirming that the correlation is robust.

To statistically validate these findings, Welch’s t-test was performed for each metric and noise level, comparing RNAtive’s ranking similarity against the null-model distribution. After applying a Benjamini-Hochberg FDR correction to the p-values, RNAtive’s performance remained significantly superior to that of a random ranker across all noise levels for every metric.

Finally, the effect size was calculated to quantify the magnitude of the performance difference. We computed Cohen’s *d* by dividing the difference in sample means by the null-model standard deviation. As shown in the lower panels of Figure 5 and Figures S8–S10 (Supplementary data), *d* remains above 1 for all metrics throughout the simulation. Given that *d >* 0.8 indicates a very strong effect, these results confirm that RNAtive is highly robust to noise in the input data.

## 4 Conclusions

RNAtive represents a novel approach for scoring and ranking ensembles of RNA 3D models predicted for a given RNA molecule. Its core innovations – the consensus-based evaluation paradigm, the fuzzy-set algorithm for interaction analysis, and the integration of 2D constraints – together provide a powerful tool for the structural bioinformatics community. RNAtive is available as a web application with an intuitive user interface, informative visualizations, and downloadable tabular data.

The tool is highly customizable through its rich set of parameters. The consensus-based approach can be extended to 2D input structures, and the workflow can incorporate additional constraints or assign weights to specific inputs, allowing users to guide the ranking generation based on their expertise. Future developments may include independent weighting of functionally important non-canonical base pairs. Different types of interactions can also be assigned distinct weights, reflecting their relative structural impact. Non-canonical interactions often play a greater role than canonical ones, although their detection can be challenging. Another promising direction is to analyze properties of the consensus structure itself. Quantifying features such as information density or similarity to top-ranked models and correlating these with ranking performance could provide a method for estimating the confidence in RNAtive-generated rankings. We believe that RNAtive will be particularly useful for experimentalists, aiding the selection of native-like RNA 3D structures from diverse *in silico* 3D predictions.

## Data and Code Availability

The source code for RNAtive and its core components is open source and available on GitHub:

- **Main application:** https://github.com/put-rnative/RNAtive
- **Bioinformatics tools wrapper:** https://github.com/tzok/cli2rest-bio
- **Interaction annotation services:** https://github.com/rnapdbee/rnapdbee-adapters

The benchmark datasets, evaluation scripts, and full results discussed in this study are openly available in a Zenodo repository at https://doi.org/10.5281/zenodo.15773391, cited as [Pielesiak et al., 2025]. The state-of-the-art tools used for comparison were run via Docker containers to ensure reproducibility, available from the repository at https://github.com/put-rnative/rna-tools-dockerized.

## Author contributions statement

MA, MS, and TZ developed the method. JP, MA, and TZ implemented RNAtive and conducted computational experiments. All authors contributed to writing and reviewing the manuscript.

## Acknowledgments

This work is supported by funds from the National Science Center, Poland [2023/51/D/ST6/01207 to TZ] and statutory funds of the Institute of Computing Science, Poznan University of Technology. We thank Bartosz Adamczyk for sharing the Dockerized version of ARES used in our benchmark analysis and Clément Bernard for insightful discussions regarding RNAtive.

