## Supplementary materials for "RNAtive to recognize native-like structure in a set of RNA 3D models"

Jan Pielesiak[1] Maciej Antczak[1,2] Marta Szachniuk[1,2]
Tomasz Zok[1]

### Supplementary Figures

The following figures provide a detailed analysis of quality assessment metrics and scoring methods on the CASP15 and randstr datasets.

**Figures S1–2: Hierarchical Clustering of Metrics** The dendrograms visualize the similarity structure among the seven quality assessment metrics used in this study. The hierarchical clustering is based on the average pairwise Spearman correlation between the metrics. The distance on the y-axis is calculated as one minus this correlation value, which groups highly correlated metrics together at lower heights.

**Figures S3–7: Ranking of Scoring Methods** The bar plots provide a comprehensive ranking of scoring methods against various reference metrics. For each scoring method (row), we report four performance measures averaged across all targets: Spearman's $\rho$, Kendall's $\tau$, the normalized Enrichment Score, and Rank-Biased Overlap. To provide a single, unified assessment, methods are ranked according to their score on the first principal component (PC1) from a Principal Component Analysis (PCA) of these four metrics. The PC1 score is labeled on each bar.

Note that an asterisk (*) indicates that for a given method, 20% or more of the underlying correlation coefficients ($\rho$ or $\tau$) were not statistically significant. Results for our method, RNAtive, tested with different parameter settings, are highlighted in bold.

---

*

[1] Institute of Computing Science, Poznan University of Technology, Piotrowo 2, 60-965, Poznan, Poland

[2] Institute of Bioorganic Chemistry, Polish Academy of Sciences, Noskowskiego 12/14, 61-704, Poznan, Poland

**Figues S8–10: Robustness Analysis** These plots assess the robustness of RNAtive's performance against increasing levels of random noise. For each ranking similarity metric, the top panel displays the mean performance and its 95% confidence interval (CI). Even at the maximum applied noise level, RNAtive's performance remains clearly distinct from that of the null model, showing no overlap between their respective 95% CIs. The bottom panel reinforces this finding, showing that the effect size – quantified by Cohen's d – remains large across the entire simulation. These results confirm that the observed performance is highly robust to noise.

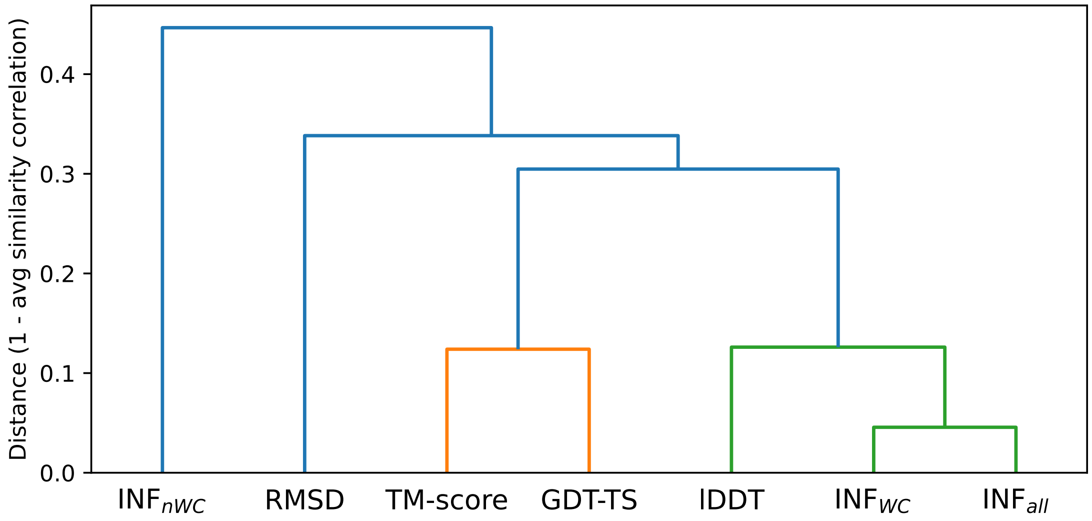


Figure S1: Hierarchical clustering of metrics for the CASP15 dataset.

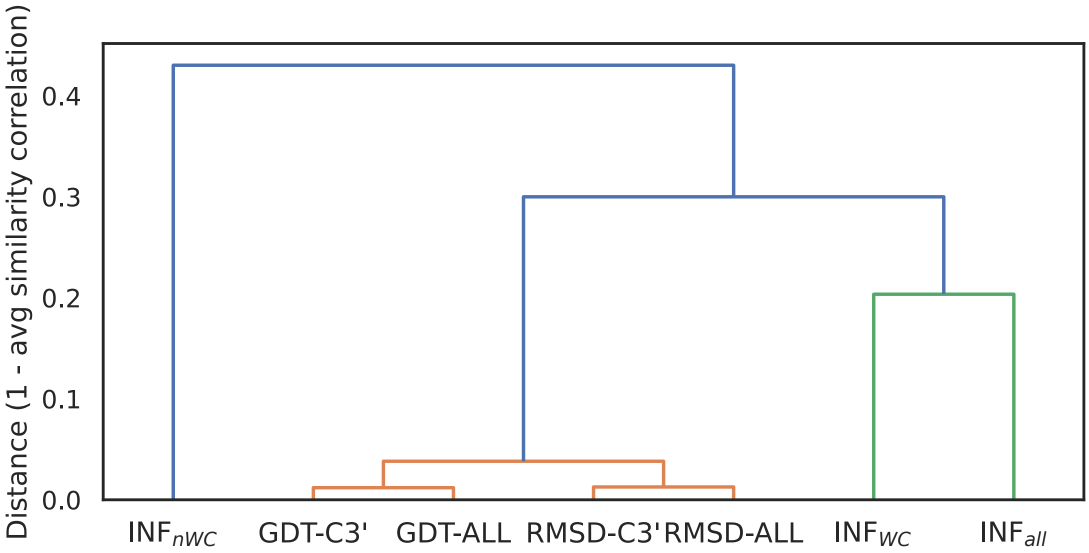


Figure S2: Hierarchical clustering of metrics for the randstr dataset.

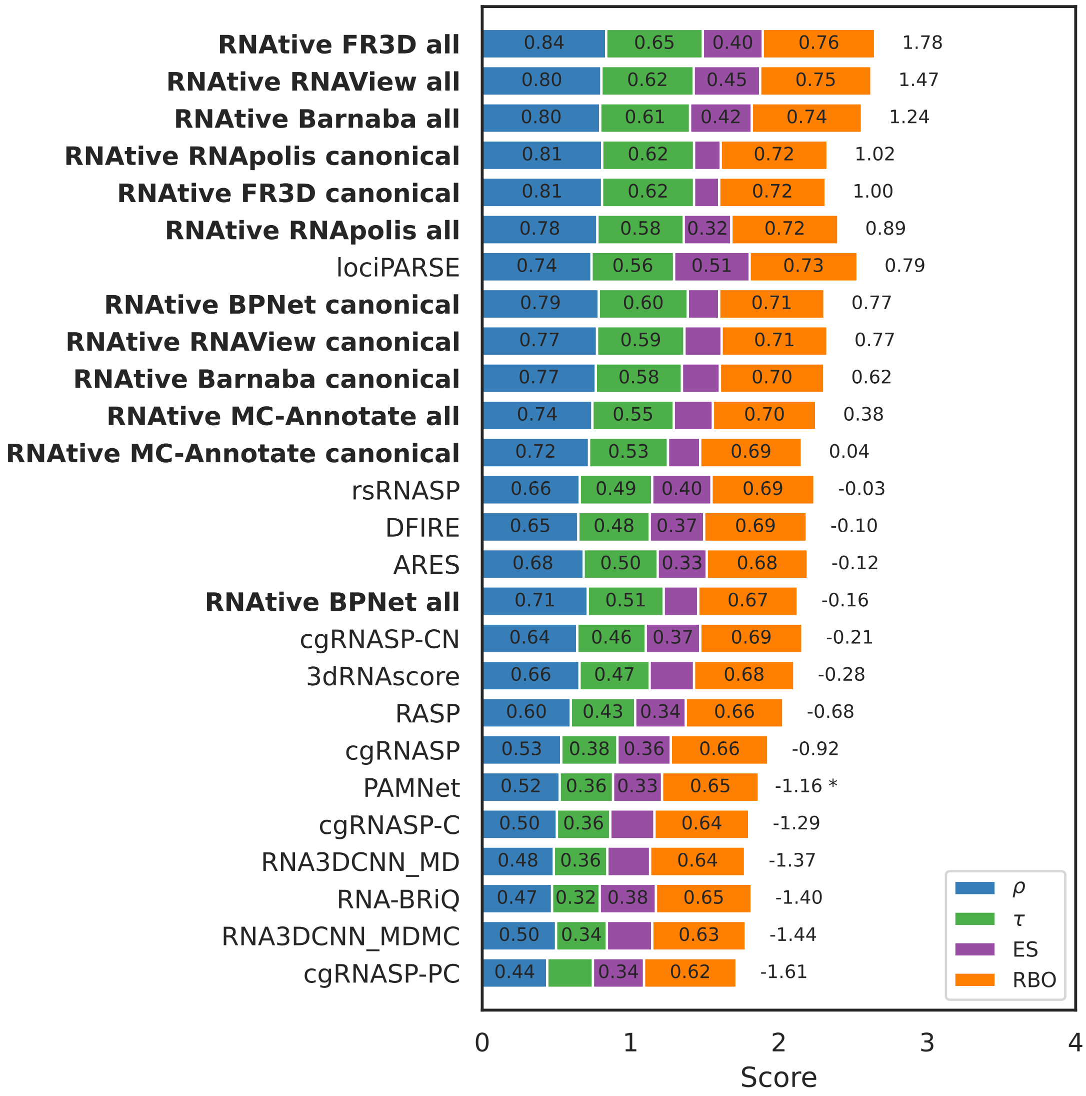


Figure S3: Ranking of scoring methods for the CASP15 dataset against the lDDT metric.

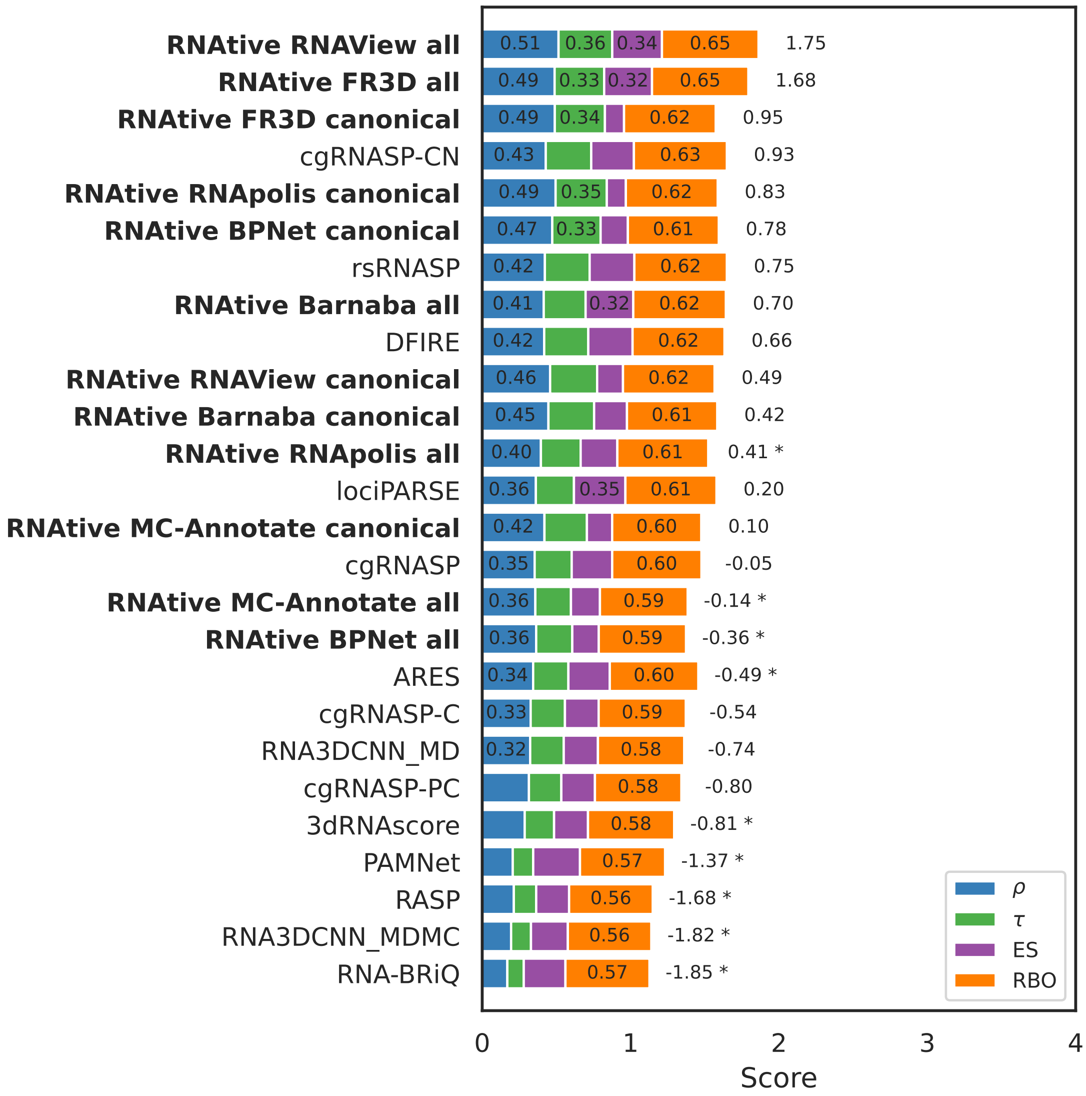


Figure S4: Ranking of scoring methods for the CASP15 dataset against the RMSD metric.

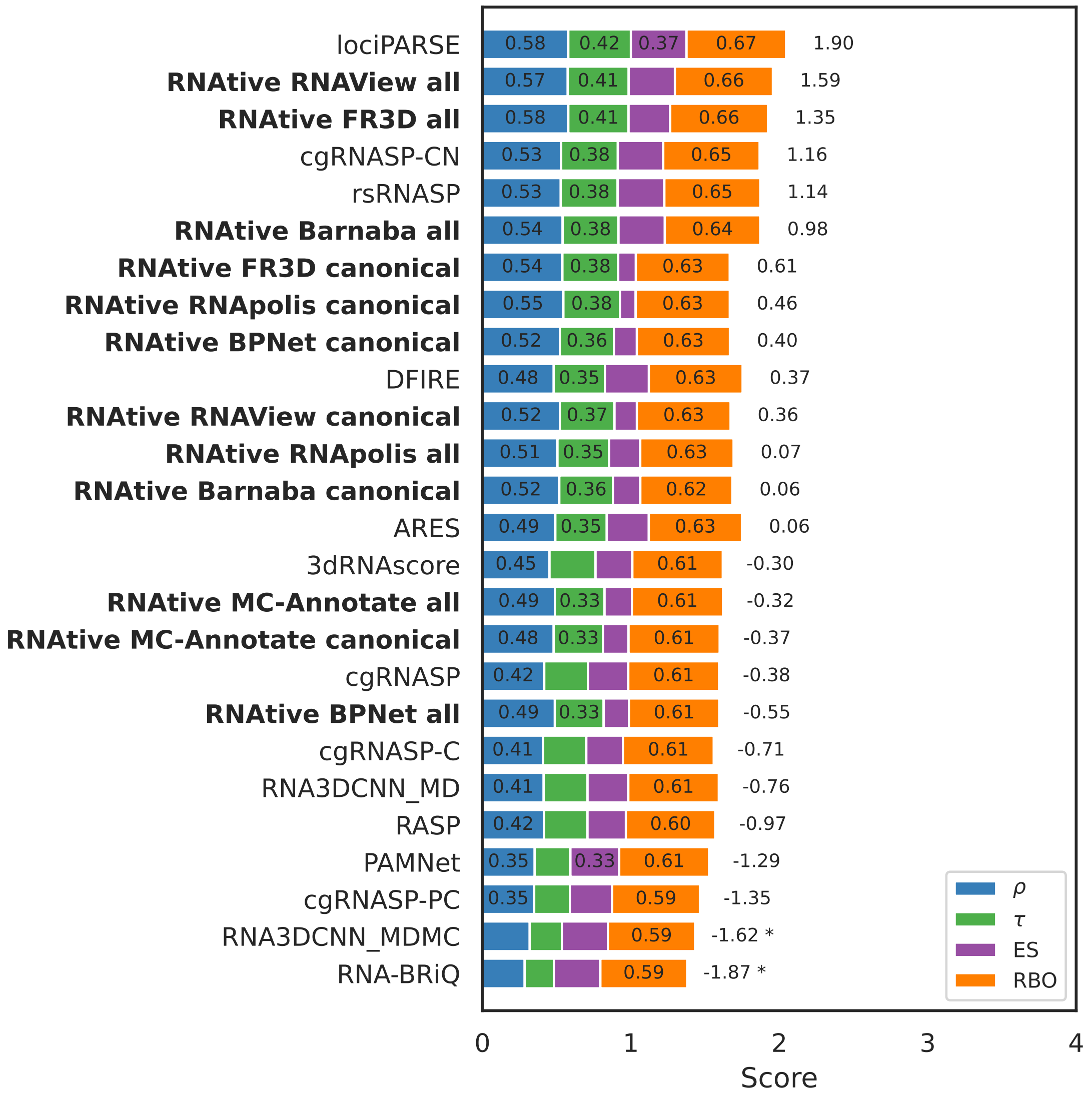


Figure S5: Ranking of scoring methods for the CASP15 dataset against the TM-score metric.

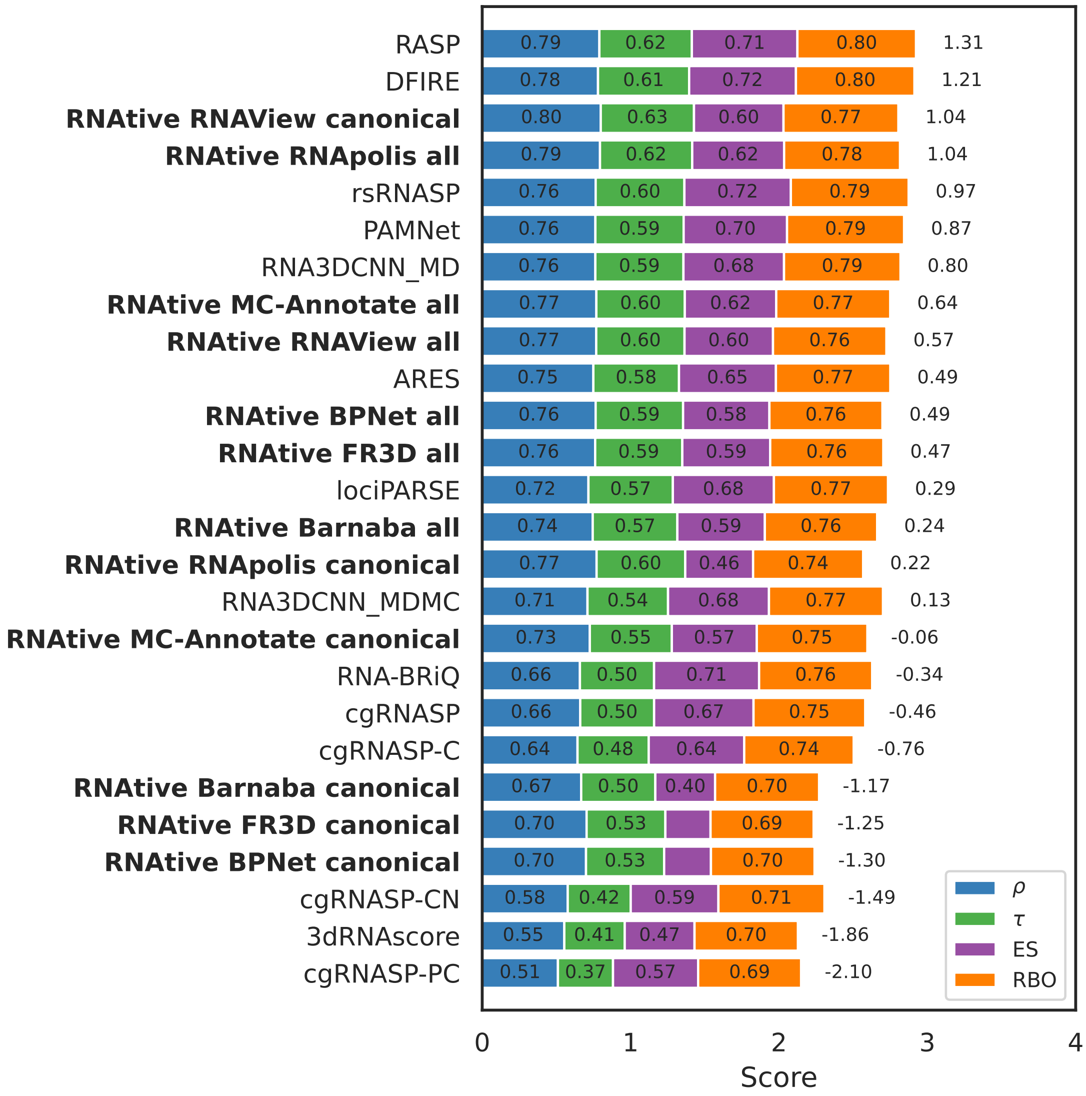


Figure S6: Ranking of scoring methods for the randstr dataset against the $\mathrm{INF}_{\mathrm{WC}}$ metric.

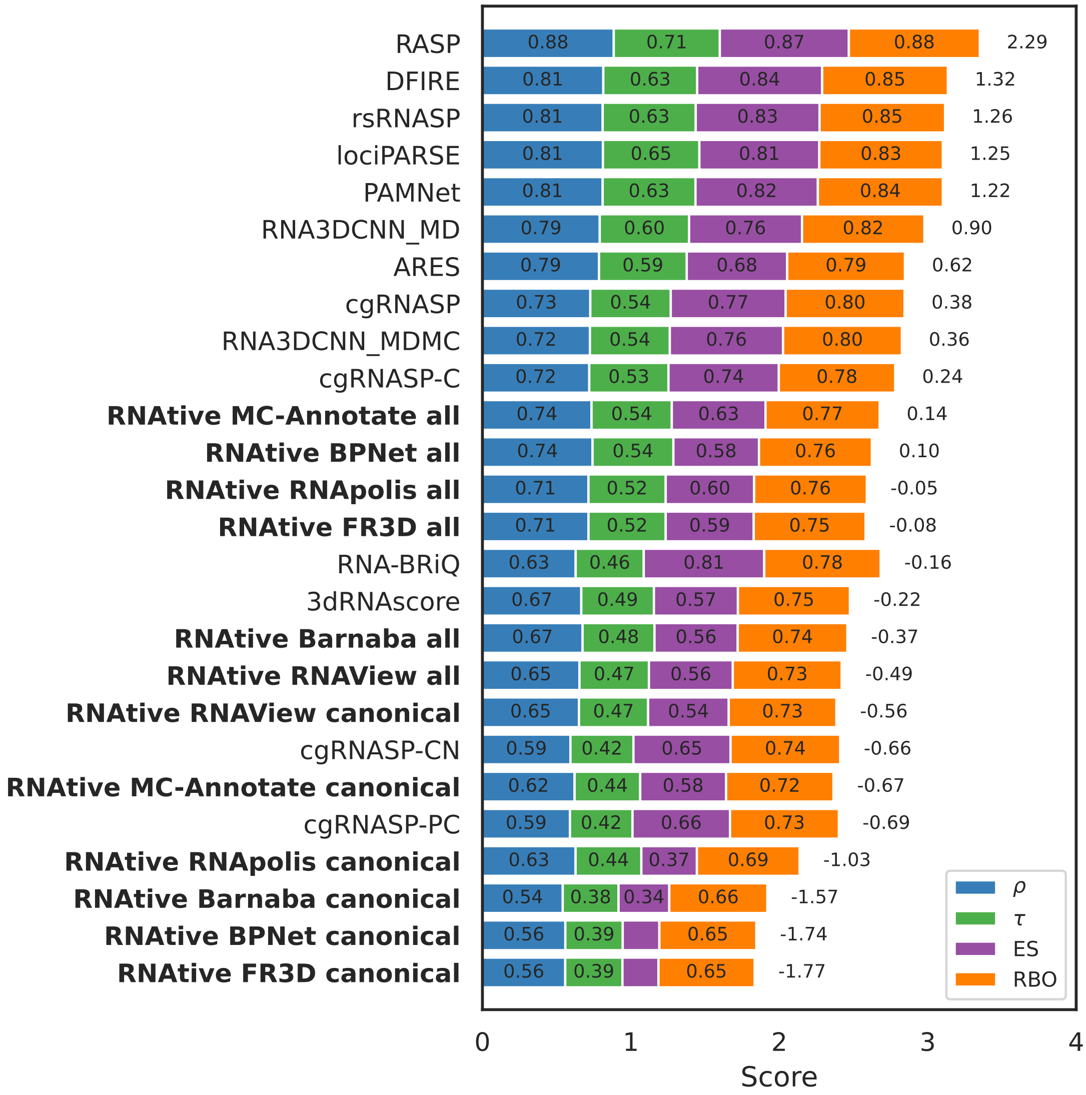


Figure S7: Ranking of scoring methods for the randstr dataset against the RMSD-ALL metric.

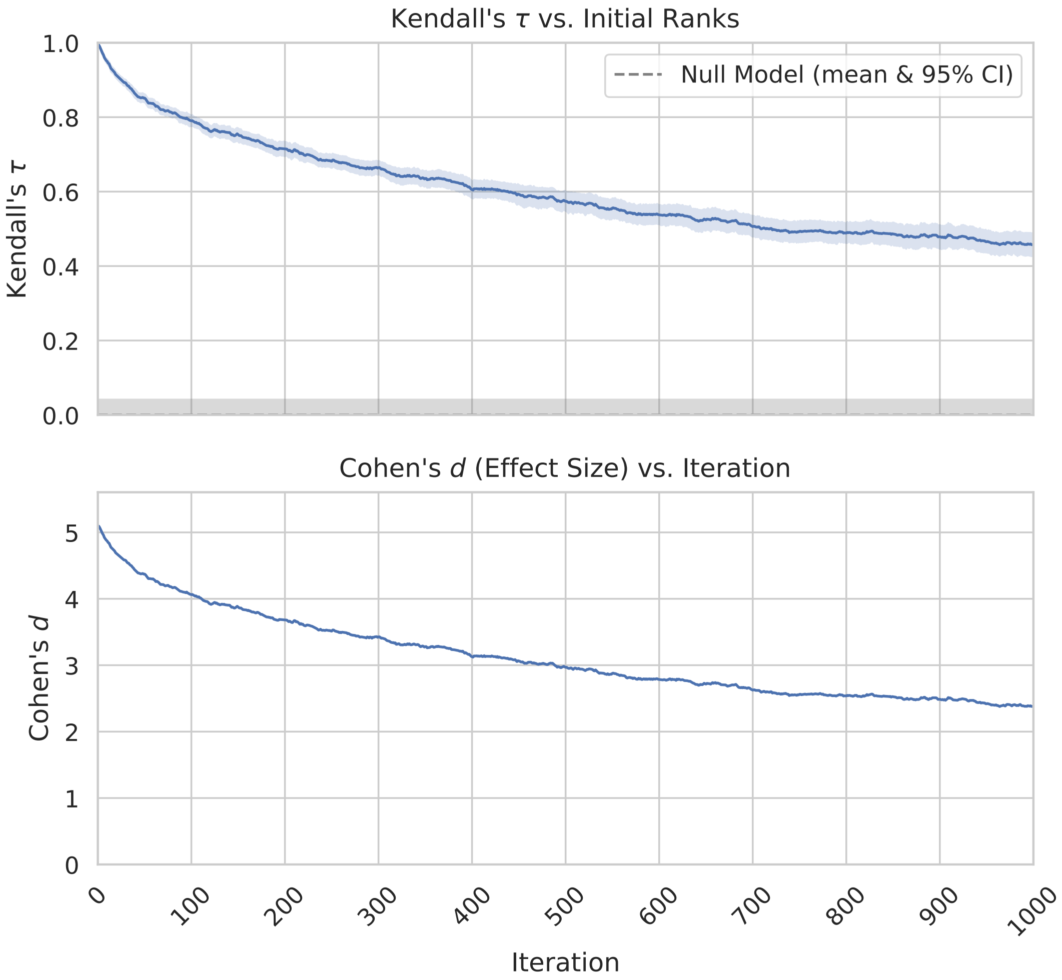


Figure S8: Robustness analysis as measured by Kendall's $\tau$.

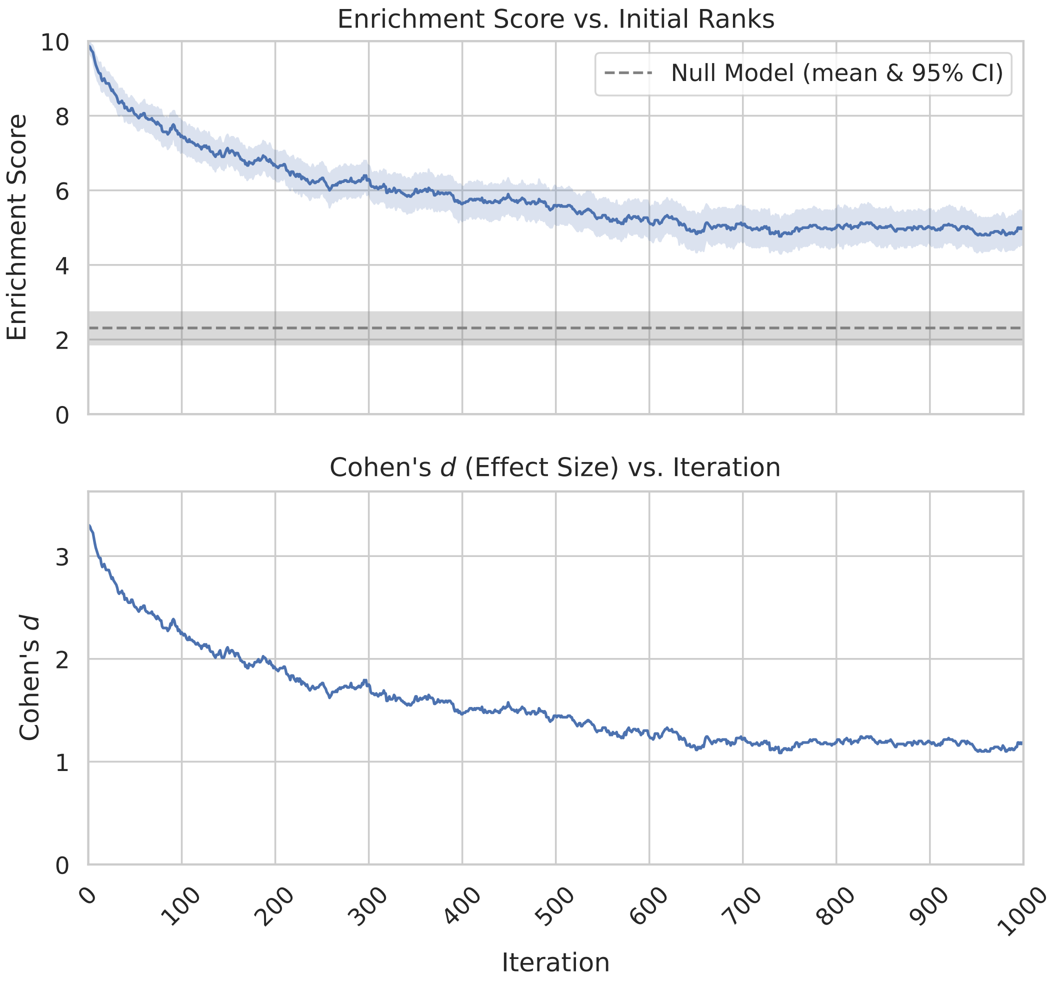


Figure S9: Robustness analysis as measured by Enrichment Score with top 25% overlap.

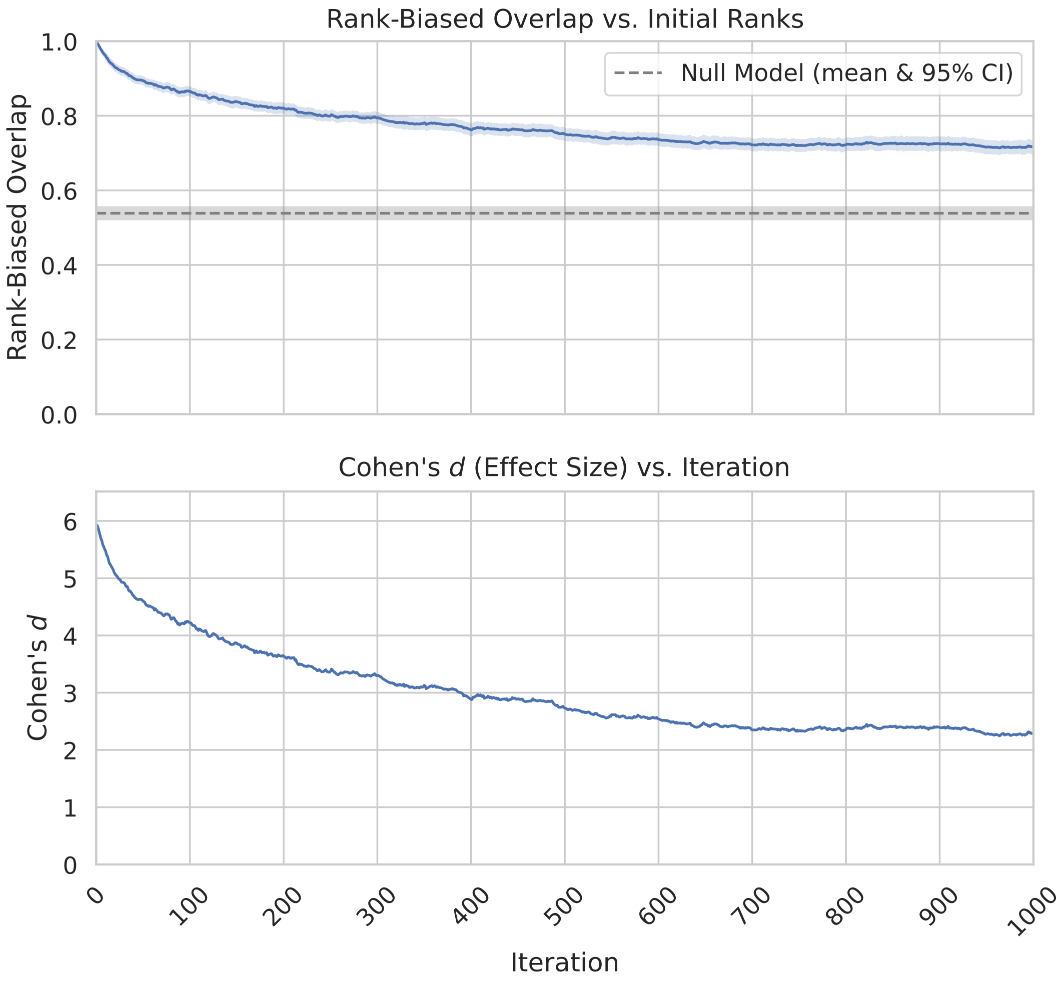


Figure S10: Robustness analysis as measured by Rank-Biased Overlap.
